# Phylogenomics of Selaginellaceae with special reference to the enigmatic *sanguinolenta* group

**DOI:** 10.1101/615633

**Authors:** Hong-Rui Zhang, Ran Wei, Qiao-Ping Xiang, Xian-Chun Zhang

## Abstract

Selaginellaceae has been repeatedly proved as monophyly by previous studies with only one genus being recognized. However, the subgeneric classification has been debated during the recent decades. Furthermore, phylogenetic position of the newly identified *sanguinolenta* group has not been resolved, varying depending on the datasets and analysis methods. We carried out the phylogenomic analyses of twenty-six species from Selaginellaceae with ten species being newly sequenced and three species representing the *sanguinolenta* group. Four of the ten newly sequenced plastomes are assembled into the complete molecules, whereas the other six species are only assembled into five to sixteen contigs owing to high numbers of repeats. The phylogenetic framework from our study is basically congruent with the subgeneric classification of Weststrand and Korall (2016b). The position of *sanguinolenta* group was resolved as the basal clade in subg. *Stachygynandrum*, which support the position β proposed by Weststrand and Korall (2016a), also supported by the morphological characters of dimorphic vegetative leaves, monomorphic sporophylls and intermixed sporangial arrangements. Both values of d*S*, d*N* and GC content in Selaginellaceae plastomes were significantly higher than those of other lycophytes (Isoetaceae and Lycopodiaceae). The correlation analysis showed that the elevated synonymous substitution rate was significantly correlated with the high GC content in Selaginellaceae. Besides, the values of d*S* and d*N* differs significantly between branches in the phylogenetic tree of Selaginellaceae. We propose that both high GC content and the extensive RNA editing sites contributed to the elevated substitution rate in Selaginellaceae, and all of these three factors could influence the stability of phylogenetic topology of Selaginellaceae.

## Introduction

Selaginellaceae, the largest family of the extant lycophytes, is one of the most ancient vascular plants with the evolutionary history dating back to the late Devonian-Early Carboniferous (370-345 Ma; (Kenrick and Crane, 1997). The Selaginellaceae recognized only one genus, *Selaginella*, with approximately 750 species (Jermy, 1990; Weststrand and Korall, 2016a; Zhou et al., 2016). Plants are herbaceous and have highly diversified growth forms, including creeping, climbing, prostrate, erect and rosetted forms, and also inhabit an impressive range of habitats, from tropical rain forests to deserts, alpine and arctic habitats (Jermy, 1990; Zhang et al., 2013).

The monophyly of the genus *Selaginella* have been repeatedly documented by previous studies, however, the subgeneric classification has been a long-term controversy during the recent decades (Korall and Kenrick, 2002, 2004; Korall et al., 1999; Weststrand and Korall, 2016a, b; Zhou et al., 2016; Zhou and Zhang, 2015). The subgeneric classification proposed by Jermy (1986) based mainly on morphological characters, such as leaf and sporophyll dimorphism and phyllotaxy has been accepted widely in the past decades. However, subsequent molecular phylogenetic analyses (Korall and Kenrick 2004, 2002; Korall, et al. 1999; Weststrand and Korall 2016a; Zhou, et al. 2016) have found the polyphyly of three out of the five subgenera: subg. *Stachygynandrum*, subg. *Heterostachys* and subg. *Ericetorum* sensu Jermy (1986). Meanwhile, a number of subclades (e.g. *sanguinolenta* group) were recovered with strong support, whereas their phylogenetic positions have not been resolved, varying depending on datasets and analyses methods used (Weststrand and Korall, 2016a; Zhou et al., 2016). For example, the *sanguinolenta* group was nested as the second basal lineage and was treated as subg. *Boreoselaginella* by Zhou et al. (2016) and Zhou and Zhang (2015), despite the conflict topologies in different analyses. More recently, Weststrand and Korall (2016a, 2016b) also recovered conflict positions for *sanguinolenta* group based on *rbcL* and two nuclear regions (*pgiC* and *gapCp*). However, they tentatively treated it as members of subg. *Stachygynandrum* with the combination of morphological evidence.

One plausible causation for the problematic phylogenetic relationship within Selaginellaceae may lie in the extreme substitution rates and the substitution rate heterogeneity of chloroplast DNA sequences among different lineages of this family (Korall and Kenrick, 2004). The branch length of the *rbcL* tree is shown unevenly distributed with more substitution sites existing in clade A (subg. *Rupestrae*, subg. *Lepidophyllae*, subg. *Gymnogynum*, subg. *Exaltatae* and subg. *Ericetorum* sensu Weststrand and Korall (2016b)) than in clade B (subg. *Stachygynandrum* sensu Weststrand and Korall (2016b)) (Korall and Kenrick, 2002). Furthermore, there is a genus-wide GC bias (above 50%) in *Selaginella* chloroplast DNA (Smith, 2009; Tsuji et al., 2007). And the presumable explanation of this GC bias is considered as a combination of reduced AT-mutation pressure and a large number of C-to-U RNA editing sites (Smith, 2009). However, whether the extreme substitution rate in Selaginellaceae is correlated with the unparalleled GC content and extensive RNA editing, and whether these factors together affect the phylogenetic positions of the unresolved lineages in Selaginellaceae remain poorly understood yet and worth for further exploration.

Nowadays, more and more recalcitrant relationships among families, lineages, or even species can be untangled using genomic (plastid, mitochondria, and nucleus) datasets, which containing unprecedented numbers of informative loci and sites, compared with multi-locus phylogeny (Lu et al., 2015; Ross et al., 2016; Zhang et al., 2016). Studies on whole plastomes of ferns and seed plants (Delsuc et al., 2005; Gao et al., 2010; Morris et al., 2018; Wei et al., 2017; Wickett et al., 2014) have been increasing quickly, whereas scarce phylogenomic studies have been carried out in lycophytes (Zhang et al., 2019). Here, for the first time, we reconstructed the robust phylogenetic framework of Selaginellaceae using plastome data from 26 species representing six out of seven subgenera based on the classification of Weststrand and Korall (2016b). We resolved the phylogenetic position of the enigmatic *sanguinolenta* group, and further discussed the correlation between phylogeny robustness, substitution rates heterogeneity and CG content bias.

## Materials and Methods

### Taxon Sampling

Thirty-six taxa (26 species of *Selaginella* and ten outgroup species) were sampled. Plastomes of ten species of Selaginella were newly sequenced; three of them were selected from *sanguinolenta* group (Table S1). Two unpublished plastomes of *S. nipponica* and *S. pallidissima* from our group (Kang et al. unpublished data) and the previously published fourteen *Selaginella* plastomes (Mower et al., 2019; Smith, 2009; Tsuji et al., 2007; Xu et al., 2018; Zhang et al., 2019; Zhang et al., 2018) were combined in the following analyses. The *Selaginella* species were selected to ensure even taxon sampling across subgenera in the genus and covered almost all major lineages (Jermy, 1990; Weststrand and Korall, 2016a, b; Zhou et al., 2016; Zhou and Zhang, 2015). All of the six subgenera sensu Zhou and Zhang (2015) were included and six out of seven subgenera sensu Weststrand and Korall (2016) were sampled in this study (Table S2). Only subg. *Exaltatae*, including two species from Africa and one species from Central and South America, was absent from this study. The outgroups include four Isoetaceae and six Lycopodiaceae species with plastomes published previously (Guo et al., 2016; Karol et al., 2010; Mower et al., 2019; Wolf et al., 2005; Zhang et al., 2017) (Table S2). GeneBank accession numbers for the sampled taxa for this study were listed in Table S2.

### DNA Extraction, Sequencing and Assembling

Total genomic DNA was isolated from silica gel dried materials with a modified cetyl trimethylammonium bromide (CTAB) method (Li et al., 2013) and sequenced on an Illumina HiSeq 2500 (Illumina, San Diego, California, USA). Plastomes were de novo assembled following the methods in Zhang et al. (2019); Zhang et al. (2018).

### Phylogenetic Analysis

Nucleotide sequences for 51 shared protein-coding genes (Table S3) from 36 plastomes were extracted and aligned individually at the protein level by MAFFT (Katoh and Standley, 2013) using the translation-aligned function in GENEIOUS v. 11.1.4 (Kearse et al., 2012). Poorly aligned regions were removed by using Gblocks v. 0.91b (Castresana, 2000) with default parameters. We generated two concatenated data matrices: (1) 51 genes and (2) 1^st^ and 2^nd^ sites of 51 genes. We further applied three partitioning strategies to the 51 gene dataset: (1) partitioning by codon sites, (2) partitioning by genes, (3) partitioning by PartitionFinder v.2.1.1 (Lanfear et al., 2017). The evolutionary model of each dataset was selected based on results from jModeltest 2.1.7 (Darriba et al., 2012).

Both maximum likelihood (ML) analyses and Bayesian inference (BI) analyses were carried out for the two datasets and ML analysis were performed for three partitioning strategies. We performed the ML analyses on the RAxML web server with 1000 bootstrap replicates and the GTR+GAMMA model (Stamatakis, 2006). We performed BI analyses in MrBayes v. 3.2.6. (Ronquist et al., 2012). Two separated MrBayes runs with four parallel Monte Carlo Markov chains (MCMC) were used, sampling every 1000^th^ generation over five million replicates, and computed a majority rule (>50%) consensus tree after removing 25% of the samples as “burn-in”. Both ML and BI trees and the supporting values are visualized using FigTree v.1.4.3 (Rambaut, 2016).

### Divergence Time Estimation

We carried out divergence time estimation to infer the time scale of the long evolutionary history within Selaginellaceae, using BEAST v. 1.8.2 (Suchard et al., 2018) under an uncorrelated lognormal relaxed clock method. Three fossil calibration nodes were employed with the first fossil calibration of the root age corresponded to the split of Lycopodiopsida and Isoetopsida (Figure 1**b**, node A: [392-451 Ma]) (Morris et al., 2018) with a selection of normal prior distributions. The second fossil calibration of the node separated Isoetaceae and Selaginellaceae (Figure 1**b**, node B: [372-392 Ma]) (Kenrick and Crane, 1997) with a lognormal prior distribution. A birth-death speciation process with a random starting tree was adopted. The MCMC chain was run for 500 million generations, sampling every 1000 generations. The effective sample size (ESS) was checked in Tracer v 1.5 (Rambaut and Drummond 2009). The maximum clade credibility tree was generated using TreeAnnotator in BEAST and the tree was plotted using FigTree v. 1.4.3 (Rambaut 2017).

**Figure 1.**
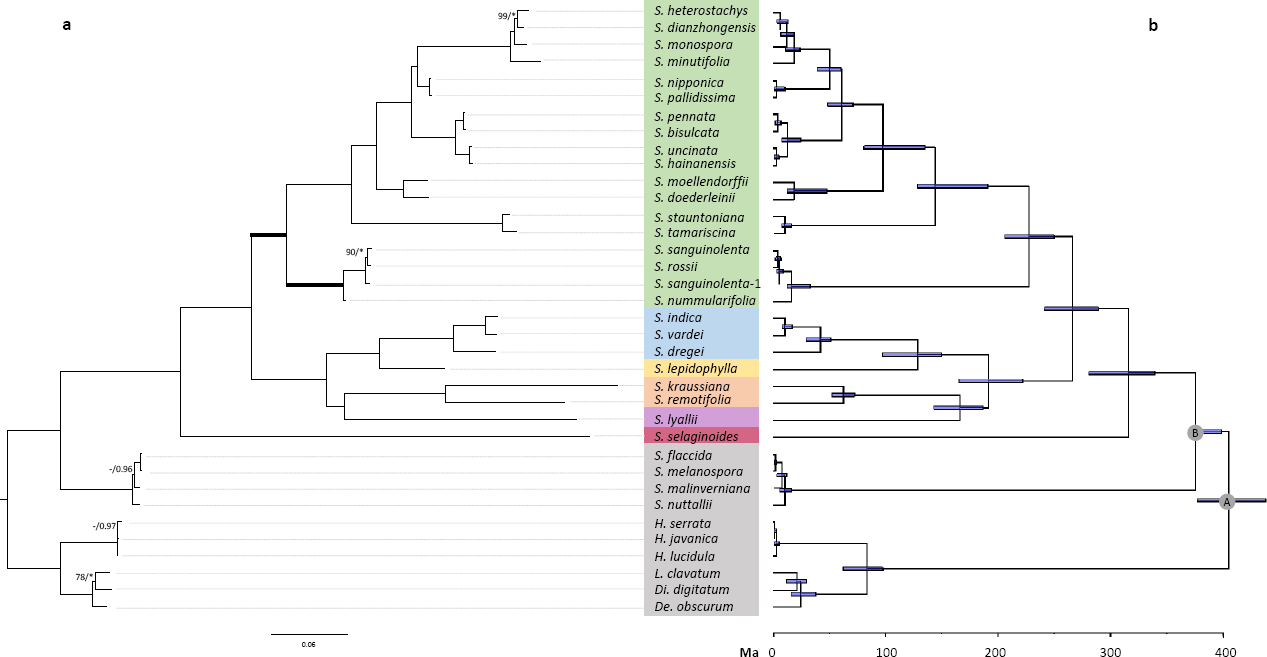
The ML phylogram of 36 species based on 51 concatenated gene-matrix and Bayesian divergence time estimation. (a) Node support values are Maximum likelihood bootstrap values (BSs)/Bayesian posterior probabilities (PPs) and are 100/1.0 unless otherwise indicated; asterisk represents 1.0 (PPs)and hyphen represents value lower than 75% (BSs); the thick lines show the resolved phylogenetic position of *sanguinolenta* group; (b) chronogram with two calibration nodes indicated by alphabets.

### Substitution Rate and GC content Analyses

The synonymous (d*S*) and nonsynonymous (d*N*) substitution rate and the ratio of d*N*/d*S* of the concatenated 51 protein-coding genes present in the plastomes of each lycophyte species were analyzed in PAML v. 4.9 (Yang, 2007), using the “branch” model (Model 1, runmode=0) implemented in the CODEML module to test if the value of d*S* and d*N* differs between branches in the phylogeny of lycophytes. In addition, the pairwise d*S* and d*N* of the concatenated 51 protein-coding genes among three lycophyte families was estimated with *Physcomitrella patens* (Sugiura et al., 2003) as reference, using PAML v. 4.9 (runmode=-2) (Yang, 2007) with codon frequencies determined by the F3 × 4 model. GC content of the shared 51 protein coding genes from the 36 plastomes were calculated in GENEIOUS 11.1.4 (Kearse et al., 2012). The difference significance of GC content and pairwise substitution rate among three families (Lycopodiaceae, Isoetaceae and Selaginellaceae) was assessed respectively using Wilcoxon rank sum tests in R v. 3.4.1 (https://www.r-project.org). The correlation between GC content and d*N*, d*S* was tested using Pearson test.

## Results

### General features of new plastomes of Selaginellaceae

We sequenced ten new plastomes of Selaginellaceae, however, only four species (*S. sanguinolenta, S. nummularifolia, S. rossii*, and *S. stauntoniana*) are assembled into the complete plastome sequence, whereas the other six species (*S. selaginoides, S. dregei, S. dianzhongensis, S. heterostachys, S. monospora* and *S. minutifolia*) are only assembled into three to sixteen contigs owing to high numbers of repeats. The plastomes of *S. sanguinolenta, S. nummularifolia, S. rossii*, and *S. stauntoniana* displayed the DR structure, which is consistent with closely related species in each lineage (*sanguinolenta* group *and S. nubigena-S. digitate* clade) *sensu* Weststrand and Korall (2016b). The length of the complete plastomes ranged from 126,835 bp (*S. stauntoniana*) to 148,924 bp (*S. nummularifolia*), whereas the total length of other six incomplete plastomes varied from ca. 117 kb to ca. 93 kb (Table 1). Remarkably, the whole set of *ndh* genes were absent from plastome of all incomplete plastomes of the six species. Only gene *ndhC* kept intact in plastome *S. stauntoniana*. Gene loss and pseudogenization occurred to a different extent in plastomes of *S. sanguinolenta, S. nummularifolia*, and *S. rossii*. The four complete plastomes contained 73-88 different genes and the gene numbers of the other five incomplete plastomes varied from 53 to 78. Overall GC content in these species ranged from 50.2% in *S. nummularifolia* to 55.3% in *S. selaginoides* (Table 1).

**Table 1.** Plastomes characteristics of newly sequenced *Selaginella* species.

| Species | Size (bp) | LSC (bp) | SSC (bp) | IR (bp) | No. different genes | No. different protein-coding genes (pseudo) | No. different tRNA genes | No. different rRNA genes | GC content (%) |
| --- | --- | --- | --- | --- | --- | --- | --- | --- | --- |
| *S. dianzhongensis* | 87,141 | - | - | - | 55 | 49 | 2 | 4 | 54.5 |
| *S. dregei* | 117,930 | - | - | - | 78 | 62 | 12 | 4 | 52.9 |
| *S. heterostachys* | 92,884 | - | - | - | 55 | 48 | 3 | 4 | 54.0 |
| *S. minutifolia* | 90,238 | - | - | - | 54 | 48 | 2 | 4 | 53.2 |
| *S. monospora* | 89,452 | - | - | - | 57 | 49 | 4 | 4 | 54.1 |
| *S. nummularifolia* | 148,924 | 53,716 | 58,186 | 18,511 | 88 | 66 (5) | 13 | 4 | 50.2 |
| *S. rossii* | 146,469 | 53,349 | 57,396 | 17,862 | 88 | 65 (6) | 13 | 4 | 50.7 |
| *S. sanguinolenta* | 148,236 | 53,606 | 57,866 | 18,382 | 86 | 65(4) | 13 | 4 | 50.9 |
| *S. selaginoides* | 84,379 | - | - | - | 53 | 42 | 7 | 4 | 55.3 |
| *S. stauntoniana* | 126,835 | 53,427 | 47,760 | 12,824 | 73 | 61 (0) | 8 | 4 | 54.1 |

### Phylogenetic relationships and evolutionary time-scale among subgenera in Selaginellaceae

Species of Selaginellaceae were recovered as monophylly for both ML and BI in all concatenated and partitioned analyses (Figure 1**b**). Within Selaginellaceae, most relationships agreed with previous phylogenetic studies with strong support values, except for the position of *sanguinolenta* group and *S. lyallii*. We found that topologies from all datasets and all partitioning strategies located *sanguinolenta* group as the basal clade in subg. *Stachygynandrum*, being sister to all other species of subg. *Stachygynandrum*, supporting the position β in Weststrand and Korall (2016a). Only topology from dataset 1 (51 genes) showed that two individuals from *S. sanguinolenta* did not form into monophyly. Besides, *S. lyallii* occupied the position as sister to subg. *Gymnogynum* with strong support value (BS 100/PP 1) with the absence of subg. *Exaltatae*.

The results of divergence time estimation showed that both Isoetaceae (ca. 10 Ma) and Lycopodiaceae (ca. 84 Ma) experienced quite recent divergence, whereas Selaginellaceae displayed a much earlier divergence time (ca. 316 Ma) and the evolution history at subgeneric level ranged from ca. 129 Ma to 227 Ma based on classification of Weststrand and Korall (2016b) (Figure 1**b**).

### Correlation between substitution rates and GC content of plastomes within Selaginellaceae and among families of lycophytes

Both values of d*S* and d*N* in Selaginellaceae plastomes were significantly higher than the other two lycophyte families (Isoetaceae and Lycopodiaceae) (P < 0.01, P < 0.01) (Figure 2 **a**). The GC content calculation of the shared 51 protein-coding genes among three families also showed that GC content is significantly higher in Selaginellaceae plastomes relative to Isoetaceae and Lycopodiaceae (P < 0.01, P < 0.01) (Figure 2 **a**). The results of correlation analysis showed that the elevated synonymous substitution rate was significantly correlated with the high GC content in Selaginellaceae (d*S*: r = 0.9596, p < 0.01; d*N*: r =0. 0.9174, P < 0.01) (Figure 2 **b**). Besides, the results using the “branch” model showed that values of d*S* and d*N* differs between branches in the phylogeny of Selaginella. The d*S* value of the branch subg. *Stachygynandrum* is extremely small, close to zero, throughout the whole phylogenetic tree, whereas the d*S* value of the other basal subgenera was quite high (Figure 3 **a**). Phylogenetic tree of d*N* displayed an opposite trend with lineages within subg. *Stachygynandrum* all showed higher value, especially the derived *Heterostachys* clade (Figure 3 **b**).

**Figure 2.**
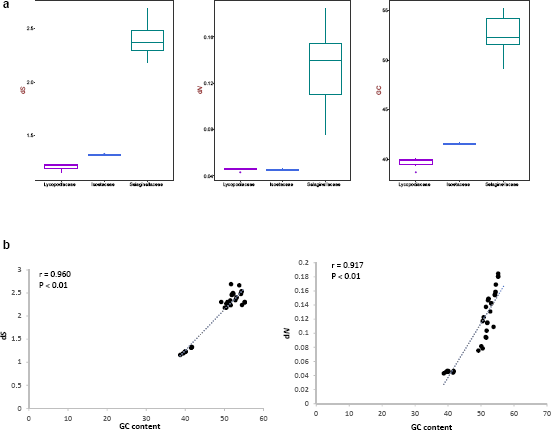
Correlation between GC content and substitution rate. (a) the value of d*S*, d*N*, and GC content in three families of lycophytes; (b) the correlation between GC content and the value of d*S*, d*N* in plastomes of lycophytes

**Figure 3.**
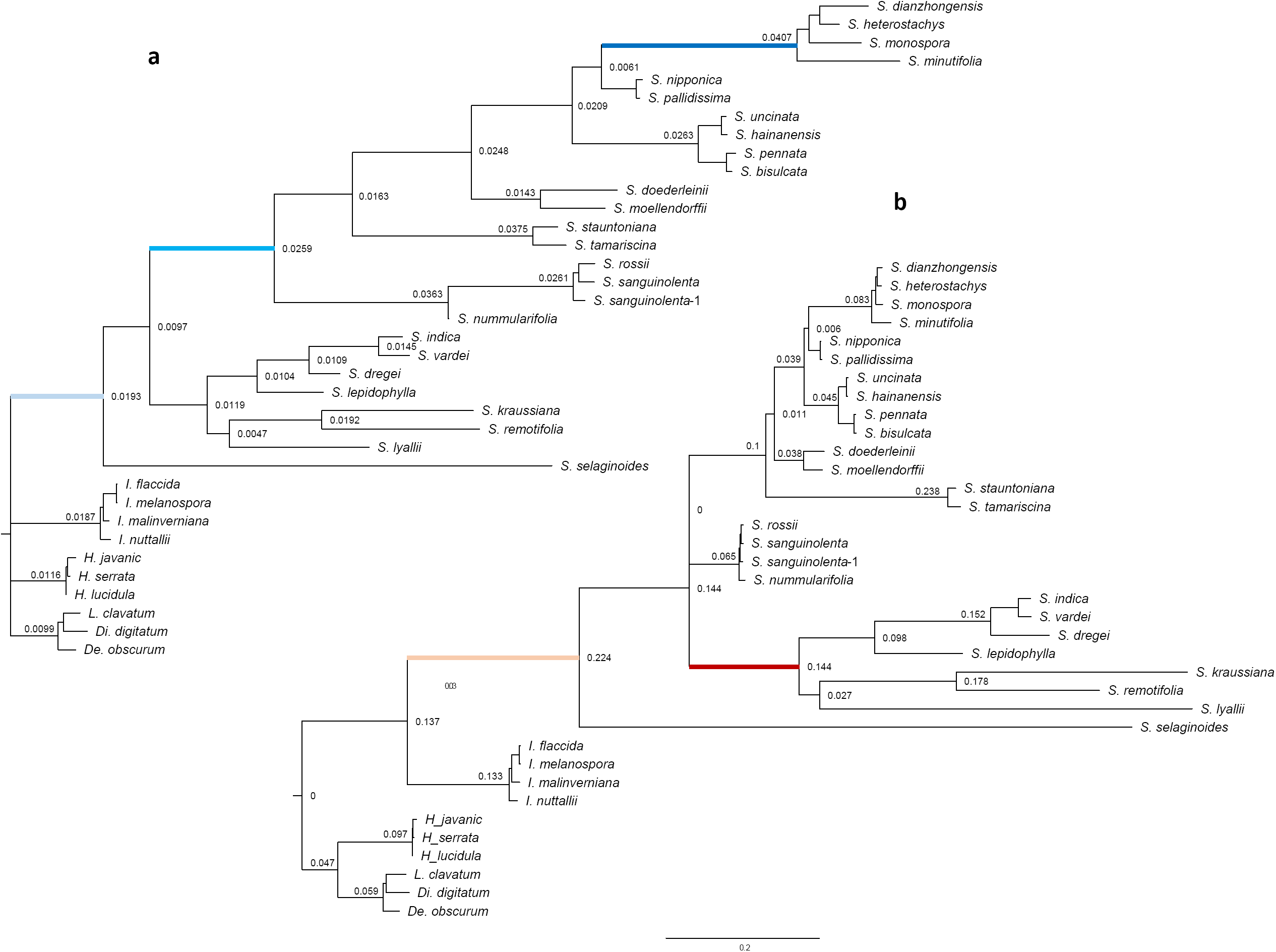
The substitution rate of different clades in Selaginellaceae based on protein-coding genes of plastomes. (a) the value of d*N* in different clades of Selaginellaceae, the light blue line represents the accelerated d*N* value in Selaginellaceae, the blue line represents the accelerated subg. *Stachygynandrum* sensu Weststrand and Korall (2016b), the dark blue line represents the accelerated *Heterostachys* group, (b): the value of d*S* in different clades of Selaginellaceae, the light orange line represents the accelerated d*S* value in Selaginellaceae, the red line represents the accelerated d*S* value in subg. *Ericetorum*, subg. *Gymnogynum*, subg. *Lepidophylla* and subg. *Rupestrae*.

## Discussion

### Phylogenetic relationships among subgenera of Selaginellaceae

The plastome phylogenies present the most robust phylogenetic framework to date of the lycophyte family Selaginellaceae. Results from all datasets and partition strategies showed highly consistent topology. The position of *sanguinolenta* group is congruent with the treatment (position β: member of subg. Stachygynandrum sensu Weststrand and Korall (2016a)) in Weststrand and Korall (2016a) from all of the analyses with strong support (BS=100/pp=1.0, Fig. 1). Only the dataset 1 showed slightly different results that two individuals from *S. sanguinolenta* are not monophyletic with *S. rossii* nested within the same clade. It is noted that these two individuals showed quite divergent growing habitat and variable morphology, collected from Xinjiang Province and Sichuan Province, respectively. We propose that the conflicting results within *S. sanguinolenta* is attributed to the inaccurate delimitation of this widely distributed species, and some putative cryptic species may exist. Further, none of our analyses recovered the phylogenetic placement of *sanguinolenta* group as the position α (sister to all *Selaginella* species except subg. *Selaginella*) in Weststrand and Korall (2016a) and the placement of Zhou et al. (2016) and Zhou and Zhang (2015). Therefore, the position α for *sanguinolenta* group is rejected. The position of *sanguinolenta* being the basal lineage in subg. *Stachygynandrum* is also consistent with morphological characters, such as dimorphic vegetative leaves arranged into four rows and monomorphic sporophylls, intermixed sporangial arrangements, and specific combination of megaspore features seen only in subg. *Stachygynandrum* (Weststrand and Korall, 2016a).

*Selaginella lyalli*i, a Madagascan spikemoss, represents subg. *Ericetorum*, one of the seven subgenera (Weststrand and Korall, 2016a). However, the relationships among subg. *Ericetorum*, subg. *Exaltatae*, and subg. *Gymnogynum* were not resolved yet (Arrigo et al., 2013; Korall and Kenrick, 2002, 2004; Korall et al., 1999; Weststrand and Korall, 2016a). The sister relationship between subg. *Ericetorum* and subg. *Gymnogynum* is well-supported from all our analyses based on different datasets and partitioning strategies. Unfortunately, the lack of representatives from subg. *Exaltatae* set up an obstacle for us to further explore the relationships among the three subgenera. However, some shared morphologies may support the unity of these three subgenera, such as the ordered, grid-like pattern in cross sections of exospores exists in subg. *Gymnogynum*, subg. *Exaltatae* and parts of subg. *Ericetorum*; and stem articulations present in species of both subg. *Gymnogynum* and subg. *Exaltatae* (Weststrand and Korall, 2016a). Therefore, we predict that subg. *Ericetorum*, subg. *Exaltatae*, and subg. *Gymnogynum* presumably clustered into one monophyletic clade, and forms sister clade to subg. *Rupestrae* and subg. *Lepidophyllae*.

### Divergence time scale of subgenera in Selaginellaceae

The result from divergence time estimation showed that the evolution of Lycopodiaceae and Isoetaceae occurred in quite recent periods (ca. 84 Ma and 10 Ma, respectively), whereas Selaginellaceae experienced extremely long evolutionary history (ca. 316 Ma) (Figure 1). Furthermore, the divergence time among subgenera also presented an unprecedently early period of differentiation. Compared with the divergence time from our results, the node age of the main lineages is basically much older than it was suggested by Klaus et al. (2017) with *sanguinolenta* group being positioned as the second diverged subgenus of Selaginellaceae. The divergence time of each subgenera in our study was largely congruent with that from Arrigo et al. (2013), except that the split time of subg. *Rupestrae* and subg. *Lepidophyllae* are much younger (ca. 129 Ma) than that of Arrigo et al. (2013) (ca. 200 Ma). Within subg. *Stachygynandrum, sanguinolenta* group diverged firstly on the phylogenetic tree (ca. 225 Ma). We presume that the extremely long evolutionary history and accumulated extensive variable DNA sites is possibly correlated with the unstable phylogenetic position.

### Substitution rate and GC content among subgenera of Selaginellaceae

Elevated substitution rate and rate heterogeneity could cause instability of phylogenetic topology, for example, of the rapid radiated lineages (Barrett et al., 2016; Wei et al., 2017; Zhang et al., 2016). The extremely high substitution rate and rate heterogeneity in Selaginellaceae have been discussed in Korall and Kenrick (2004), and the phylogenetic tree have shown that branch length of clade A is relatively longer than that of clade B, indicating more site substitutions have occurred in clade A (Korall and Kenrick, 2004). The results of the synonymous (d*S*) and nonsynonymous (d*N*) substitution rate for each lineage of Selaginellaceae showed that the branch representing subg. *Selaginella* possess quite high value for both d*N* and d*S*, owing to the early split from other subgenera in Selaginellaceae. The correlation between substitution rate and generation time (GT) has been shown in previous studies that species with faster generation time tend to have higher substitution rates due to the more frequent genome copy and more DNA replication errors per unit time (Lehtonen and Lanfear, 2014; Thomas et al., 2010). The generation time of most Selaginellaceae species are annual, and therefore, the early divergence time of subg. *Selaginella* may result in higher substitution rate. The branch length of clade A and clade B is also unevenly distributed for both d*N* and d*S*. However, the branch length of clade A is relatively shorter than that of clade B in the d*N* tree, whereas the branch length of clade A is basically longer than that of clade B in the d*S* tree. The higher value of d*N* for clade B reflected that positive selective took place for survival and adaptation to the diverse habitats after the differentiation from clade A (Gillespie, 1991; Yang and Nielsen, 2000). The higher value of dS for clade A reflected the extremely long evolutionary history and accumulated synonymous substitution sites, compared with the younger subg. *Stachygynandrum* (Thomas et al., 2010).

The pairwise d*N*, d*S* value and GC content of the concatenated 51 protein-coding genes presented significantly high in Selaginellaceae. Furthermore, the correlation analyses between substitution rate (d*N*, d*S*) and GC content showed strong positive correlation (d*S*: r = 0.9596, p < 0.01; d*N*: r =0. 0.9174, P < 0.01). The unparalleled high GC content not only exists in plastomes but also in mitochondrial genomes, in which the extremely high level of RNA editing has been both reported (Hecht et al., 2011; Oldenkott et al., 2014). Therefore, RNA editing is proposed to be tightly connected to the high GC content of the Selaginellaceae organelle genomes (Mower et al., 2019; Smith, 2009). In summary, we propose that both high GC content and extensive RNA editing contribute at least partly to the elevated substitution rate in Selaginellaceae, and all three factors could have a compound impact on the stability of phylogenetic reconstruction of Selaginellaceae.

## Acknowledgments

We thank Bertier J. C., Liu B., Zhao C. F. and Zhu Y. M. for help with materials collection. This work was supported by the National Natural Science Foundation of China (grant number 31670205, 31770237).

## Supplementary materials

Table S1 Voucher information of newly sequenced species of Selaginellaceae in this study.

Table S2 Taxonomic distribution and GenBank accession numbers for the taxa included in the study.

Table S3 List of fifty-one protein-coding genes included in the phylogenetic analysis.

