## Supplementary table 1 and 2 for "Phylogenomics of Selaginellaceae with special reference to the enigmatic *sanguinolenta* group"

**Table S1** Voucher information of newly sequenced species of Selaginellaceae in this study

| Species | Collection Number | Collector | Locality | Herbarium |
|---|---|---|---|---|
| *S. dianzhongensis* | 8158 | Y.-M. Zhu | Yimen, Yunnan | PE |
| *S. dregei* | 26698 | B. Liu | Kenya | PE |
| *S. heterostachys* | 83 | H.-R. Zhang | Tianquan, Sichuan | PE |
| *S. minutifolia* | 7764 | X.-C. Zhang et al. | Jinping, Yunnan | PE |
| *S. monospora* | 7770 | X.-C. Zhang et al. | Jinping, Yunnan | PE |
| *S. nummularifolia* | 9047 | C.-F. Zhao | Lasha, Tibet | PE |
| *S. rossii* | 199 | H.-R. Zhang | Korea, Incheon | PE |
| *S. sanguinolenta*-1 | 8829 | X.-C. Zhang | Jiulong, Sichuan | PE |
| *S. selaginoides* | Tour09165 | Bertier J. C. | France, Chamonix | PE |
| *S. stauntoniana* | 7644 | X.-C. Zhang | Huairon, Beijing | PE |

**Table S2** Taxonomic distribution and GenBank accession numbers for the taxa included in the study.

| **Family** | **Subgenera 2015** | **Subgenera 2016** | **Species** | **GenBank accession number** |
|---|---|---|---|---|
| Selaginellaceae | *Heterostachys* | *Stachygynandrum* | *S. uncinata* | MG272483 |
| | | | *S. hainanensis* | MH598533 |
| | | | *S. bisulcata* | MH598531 |
| | | | *S. pennata* | MH598534 |
| | | | *S. nipponica* | - |
| | | | *S. pallidissima* | - |
| | | | *S. heterostachys* | - |
| | | | *S. monospora* | - |
| | | | *S. proniflora* | - |
| | | | *S. dianzhongensis* | - |
| | *Stachygynandrum* | | *S. moellendorffii* | MG272484 |
| | | | *S. doederleinii* | MH598532 |
| | *Pulviniella* | | *S. tamariscina* | MH598537 |
| | | | *S. stauntoniana* | MK622384 |
| | *Boreoselaginella* | | *S. sanguinolenta* | MH598536 |
| | | | *S. sanguinolenta*_1 | MK622383 |
| | | | *S. rossii* | MK622382 |
| | | | *S. nummularifolia* | MK622381 |
| | *Ericetorum* | *Ericetorum* | *S. lyallii* | MK156800 |
| | | *Gymnogynum* | *S. remotifolia* | MH598535 |
| | | | *S. kraussiana* | MH549643 |
| | | *Rupestrae* | *S. vardei* | MG272482 |
| | | | *S. indica* | MK156801 |
| | | | *S. dregei* | - |
| | | *Lepidophyllae* | *S. lepidophylla* | MK089531 |
| | *Selaginella* | *Selaginella* | *S. selaginoides* | - |
| | | | *I. flaccida* | NC 014675 |
| | | | *I. malinverniana* | MH549640 |
| | | | *I. melanospora* | NC 038072 |
| | | | *I. nuttallii* | NC 038073 |
| Lycopodiaceae | | | *H. javanica* | KY609860 |
| | | | *H. lucidula* | NC 006861 |
| | | | *H. serrata* | NC 033874 |
| | | | *De. obscurum* | MH549637 |
| | | | *Di. digitatum* | MH549638 |
| | | | *L. clavatum* | MH549642 |
| Outgroup | | | *Physcomitrella patens* | AP005672 |

**Table S3** List of fifty-one protein-coding genes included in the phylogenetic analysis

| **Category** | **Genes** |
|---|---|
| Photosystem I | *psaA*, *psaB*, *psaC*, *psaI*, *psaJ* |
| Photosystem II | *psbA*, *psbB*, *psbC*, *psbD*, *psbE*, *psbF*, *psbH*, *psbI*, *psbJ*, *psbK*, *psbL*, *psbM*, *psbN*, *psbT*, *psbZ* |
| Cytochrome b6/f complex | *petA*, *petB*, *petD*, *petG*, *petL*, *petN* |
| ATP synthase | *atpA*, *atpB*, *atpE*, *atpF*, *atpH*, *atpI*, |
| ribosomal protein | *rpl2, rpl16, rpl22, rps3, rps4, rps7, rps8, rps11, rps14, rps18, rps19* |
| RNA polymerase | *rpoA, rpoB, rpoC1* |
| Other genes | *ccsA, clpP, rbcL, ycf3, ycf12* |
